# Biomechanical characteristics of scapula and glenohumeral movements during pitching motion in injury-prone college baseball pitchers

**DOI:** 10.1101/2021.11.22.469507

**Authors:** Koji Miyashita, Sentaro Koshida, Taro Koyama, Kenicihro Ota, Yusuke Tani, Ryoji Okamune

**Affiliations:** Department of Physical Therapy, College of Life and Health Sciences, Chubu University, Matsumoto-cho 1200, Kasugai City, Aichi, Japan 487-8501; Department of Judotherapy and Sports Medicine, Faculty of Health Sciences, Ryotokuji University, 5-8-1 Akemi, Urayasu, Chiba, Japan 279-8567; Matsushita Orthopedics, Matsumoto-cho 1200, Kasugai City, Aichi, Japan 487-8501; Watanabe Orthopedics and Rehabilitation Clinic, Matsumoto-cho 1200, Kasugai City, Aichi, Japan 487-8501; Advanced Reha Co.ltd, Matsumoto-cho 1200, Kasugai City, Aichi, Japan 487-8501

## Abstract

Coordination of glenohumeral and scapular movements plays an important role in the injury prevention of baseball pitchers. However, there is no objective data establishing the direct relationship between pitching injuries and associated glenohumeral and scapular movements. Therefore, the objectives of the present study were to demonstrate biomechanical differences in scapular and glenohumeral movements during pitching between injury-prone pitchers and healthy college baseball pitchers. Thirty collegiate baseball pitchers were classified into two groups according to their injury status: injury-prone group (N=15, 20.7±1.4 years, 180.1±6.5 cm, 78.9±5.4 kg) and control group (N=15, 20.9±1.1 years, 177.1±6.6 cm, 72.3±6.7 kg). We obtained the pitching motion data using the three-dimensional motion analysis technique with four high-speed cameras. The horizontal abduction angles of the glenohumeral joint during cocking and acceleration phases were significantly greater in injury-prone pitchers [19.0° (95% CI: 14.4–23.6) at foot contact, −4.0° (95% CI: −7.7 to −0.2) at maximum external rotation (MER), and −0.3° (95% CI: −4.8 to −4.2) at ball release] than in healthy controls [11.7 °(95%CI:7.1 to 16.3) at foot contact, −10.0°(95%CI: −13.7 to −6.3) at MER, and −6.9°(95%CI: −11.4 to −2.4)](*p*<0.01). In addition, the external rotation angle (ER) of the scapula at MER was significantly greater in the injury-prone group [−0.1° (95% CI: −5.0 to 4.8)] than in the control group [−12.3° (95% CI: −17.2 to −7.4)] (p<0.01), but there was no difference in the scapular ER during foot contact between the two groups. These results suggests that injury-prone pitchers have less internal rotation of the scapula and more horizontal abduction of the glenohumeral joint during cocking and acceleration phases. Therefore, sports medicine practitioners may need to pay more attention to coordination of scapular and glenohumeral movements during the cocking and acceleration phases of pitching for prevention of shoulder injuries.

## Introduction

The scapula plays a pivotal role in the kinetic chain, transferring energy derived from trunk rotation to the pitching arm [1,2]. Therefore, scapular dysfunction is believed to lead baseball pitchers to injuries, often occurring in the glenohumeral joint [3,4]. Despite evidence suggesting a link between scapula dysfunction and shoulder pain, however, there is no consensus on a direct relationship between both scapular malposition/malorientation and pitching injuries [5]. Burkhart et al. [6] proposed the concept of SICK (scapular malposition, inferior medial border prominence, coracoid pain and malposition, and dyskinesis of scapula movement) to explain the scapular asymmetry seen in injured overhead athletes. However, several studies have reported that overhead athletes have distinctive characteristics in the scapular position and orientation regardless of shoulder injury [5,7–11]. These results suggest that the SICK scapula may be an adaptive change for efficient pitching performance, rather than a pathological change leading to pitching-related injuries. Scapular dysfunction in pitching motion may be more reliable predictor of pitching-related shoulder injuries. The concept of hyperangulation [12] suggests that scapula restriction during the cocking phase increases the horizontal abduction of the humerus relative to the glenoid fossa and increases the mechanical stress placed on the shoulder at maximum external rotation (MER) and abduction [12,13]. It has been clinically recognized that this dynamic malalignment of scapular-glenohumeral complex is associated with pitching-related shoulder injuries [14,15]. Therefore, the purpose of this study was to individually quantify scapular and glenohumeral joint movements during baseball pitching and to compare biomechanical differences between the groups of baseball pitchers who prone to shoulder injuries and their counterparts. We hypothesized that injury-prone college baseball pitchers would have less scapular motion and more glenohumeral motion when throwing compared to healthy controls.

## Materials and methods

### Participants

The study participants were 30 male college baseball pitchers who pitch in an overhand style. They were members of a college baseball team and had participated in games for four years, somewhere between 2012 and 2019. One team athletic trainer provided conditioning and injury prevention programs for these players, and kept daily records of their complaints and conditions. Based on these records, the participants were classified into two groups of 15 participants each according to the injury. The injury-prone group consisted of 14 right-handed and one left-handed pitcher who had sustained repeated non-time-loss shoulder injuries related to pitching within the past three months. A non-time-loss injury is defined as injury that does not result in restriction from participation for at least 24 hours (not restriction from participation beyond the day of injury), and this definition has been used for injury research in sports [16–18]. We chose the non-time-loss criterion because the severe pain in the throwing shoulder that causes a time-loss injury may alter the original pitching motion. We also excluded pitchers who complained of upper extremity pain during the measurement. The control group consisted of 12 right-handed and 3 left-handed pitchers who had no shoulder pain when pitching within the past 6 months. A signed informed consent was obtained from each participant prior to participation in the study. The study protocol was approved by the Ethics Committee of Chubu University, Aichi, Japan. Approval Number: 270008.

### Pitching motion analysis

Pitching motion data were collected in an indoor biomechanics laboratory using the procedure previously [19]. The pitcher’s plate was set up 18 m away from a fixed target (3.0 cm diameter) placed in the center of a net set up to catch the ball behind a home base. Each participant wore a baseball glove on the non-pitching side and tight-fitting spandex shorts, socks, and shoes with spikes.

Fig 1 (left) shows the marker placement in this study. The reflective markers (1.0 cm diameter) were placed at bony landmarks of the participant as follows: the spinous processes of the seventh cervical (C7), the eighth thoracic (Th8), and the first lumbar (L1) vertebrae, and the manubrium of sternum. In addition, lightweight urethane bars (20 cm [length] × 1 cm [width] × 1 cm [height]) with reflective markers attached to the both ends (Fig 1, inset) were placed on the acromion process, the dorsal side of the distal end of the humerus, and the distal end of the forearm on the pitching side. The acromion bar marker was placed on the flat part of the acromion process so that the two reflective markers affixed on both ends were aligned from to back. Therefore, the movement of the acromion bar markers indicated the anterior–posterior tilt and external–internal rotation of the scapula. In addition, two other bar markers were placed on the humerus and the forearm to mark the perpendicular direction.

**Fig 1.**
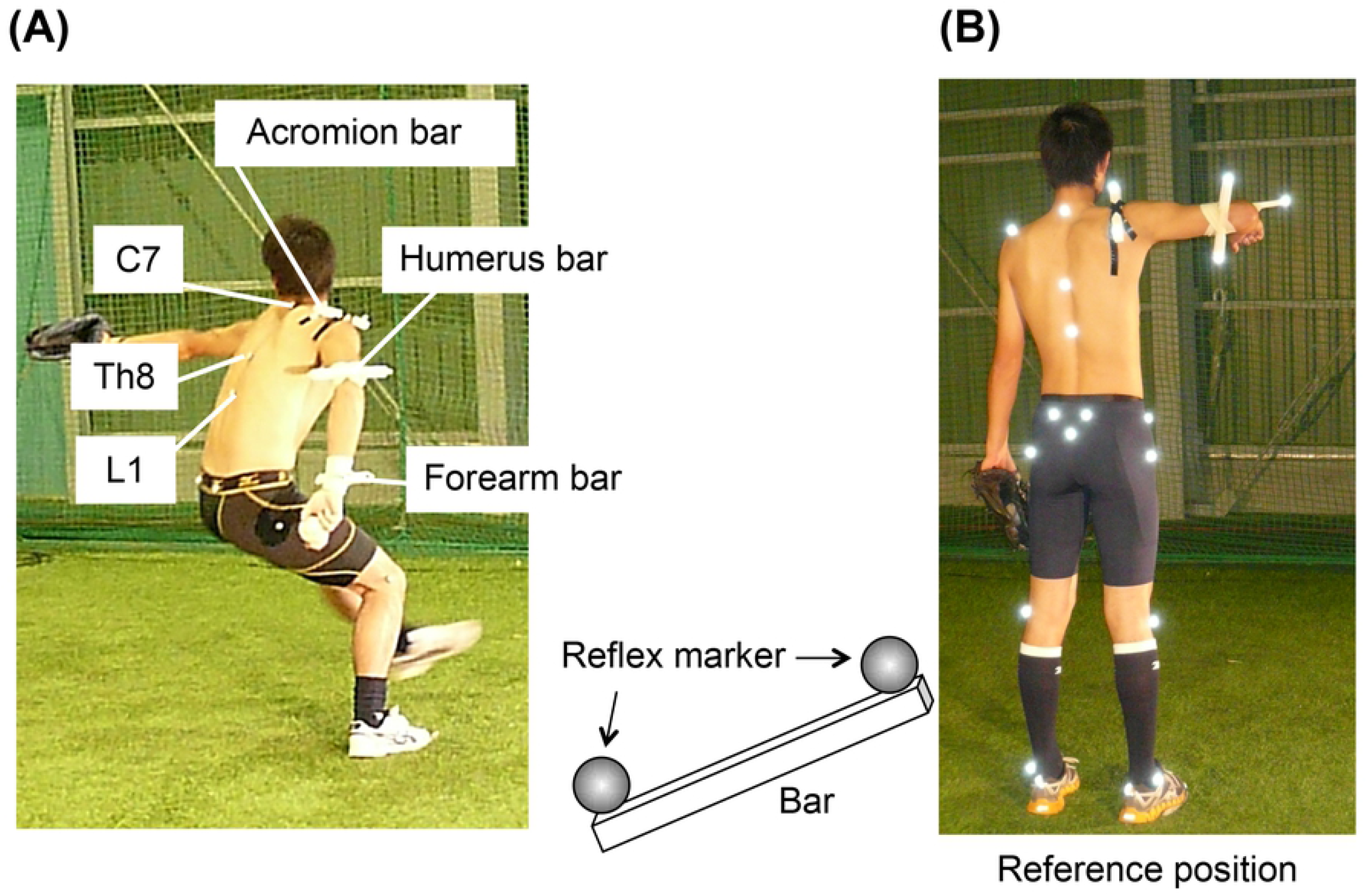
Points for the marker placement and the reference position. (A) The reflective markers were placed at bony landmarks of the participant as follows: the spinous processes of the seventh cervical (C7), the eighth thoracic (Th8), and the first lumbar (L1) vertebrae, and the manubrium of sternum. lightweight urethane bars with reflective markers were affixed on both of the edges, on the acromion process, the dorsal side of the distal end of the humerus, and the distal end of the throwing forearm., (B) B shows reference position, shoulder is 90 deg flexion and abduction, elbow is 90 deg flexion, and forearm is neutral position.

After a normal warm-up, each participant made 10 pitches toward the target with maximum effort. The pitching motion data were obtained using four high-speed cameras (IEEE1394b high-speed camera FKN-HC200C, 4 Assist, Tokyo, Japan), and four video images at 200 frames per second were electrically synchronized. The four cameras were placed at the right and left rear, at the dominant hand side, and in front of the participant. We edited the motion video frame by frame as shown in the attached file “video1.”

In this study, the most accurate throw to a fixed target among the pitching attempts of each participant was selected for analysis. After transferring the motion data to a personal computer (ESPRIMO WD2/A3, Fujitsu, Tokyo, Japan), the video images were superimposed on a computer display, and the markers were automatically tracked using a 2- to 3-dimensional motion analyzer (Frame-DIAS V, DKH Inc., Tokyo, Japan). Then the digitized points were obtained by using direct linear transformation procedures [20]. The low-pass filtering was performed at 15 Hz. The computed data were external rotation (ER) angle of the shoulder complex, the ER angle of the glenohumeral joint, the posterior tilting angle of the scapula, the horizontal abduction angle of the shoulder complex, horizontal abduction of glenohumeral joint, and the ER of the scapula from the foot contact to the ball release.

Fig 2 illustrates the kinematic models used in the study [19]. To calculate the shoulder ER, glenohumeral ER, and posterior scapular tilt angles, two corresponding triangles were set up between the markers to define the body segments. For the calculation of the shoulder ER angle, one triangle was formed between the L1 marker and each midpoint of the acromion bar and humeral bar markers, and another triangle was formed between midpoints between each the forearm marker and the humeral bar and acromion bar markers. Similarly, for the calculation of the gleohumeral ER angle, one triangle was formed between the front and rear markers of the acromion bar and the rear marker of the humeral bar, and another triangle was formed between the front and rear markers of the humeral bar and the rear marker of the acromion bar. Finally, to calculate the posterior tilt angle of the scapula, one triangle was formed between the C7 and Th8 markers and the midpoint of the acromion bar, and another triangle was formed between the midpoint of the acromion bar and the posterior marker and the C7 marker. Once the triangles are formed, the unit vector of the normal projected from each corresponding triangle and the inner product of the two unit vectors are calculated. The cosine angle of that inner product was used as each joint angle. For the calculation of the glenohumeral horizontal abduction angle and the scapular ER angle, one unit vector of the normal direction projected from the triangle connecting the C7 marker, the Th8 marker, and the median sternal marker was calculated. The scapular ER angle was calculated by the inner product of this unit vector in the normal direction and the acromion bar. The glenohumeral horizontal abduction angle was defined as the angle in the horizontal plane between the acromion bar and the line connecting the midpoint of the acromion bar and the midpoint of the humeral bar. The cosine angle of the inner product was used as each joint angle. Posterior scapular tilt angle, glenohumeral ER, scapular ER, and horizontal scapulohumeral abduction angle were set to 0° when the participant stood upright with shoulder abduction and elbow flexion of 90° (Fig 1, right). All kinematic data were normalized on a 100% scale from the time of foot contact in the stride to the time of ball release to facilitate comparison between participants. As a preliminary study, the author verified the validity of the measurements of the posterior tilt angle of the scapula during external rotation of the shoulder joint used in this study by comparing them with data obtained by electromagnetic sensors.

**Fig 2.**
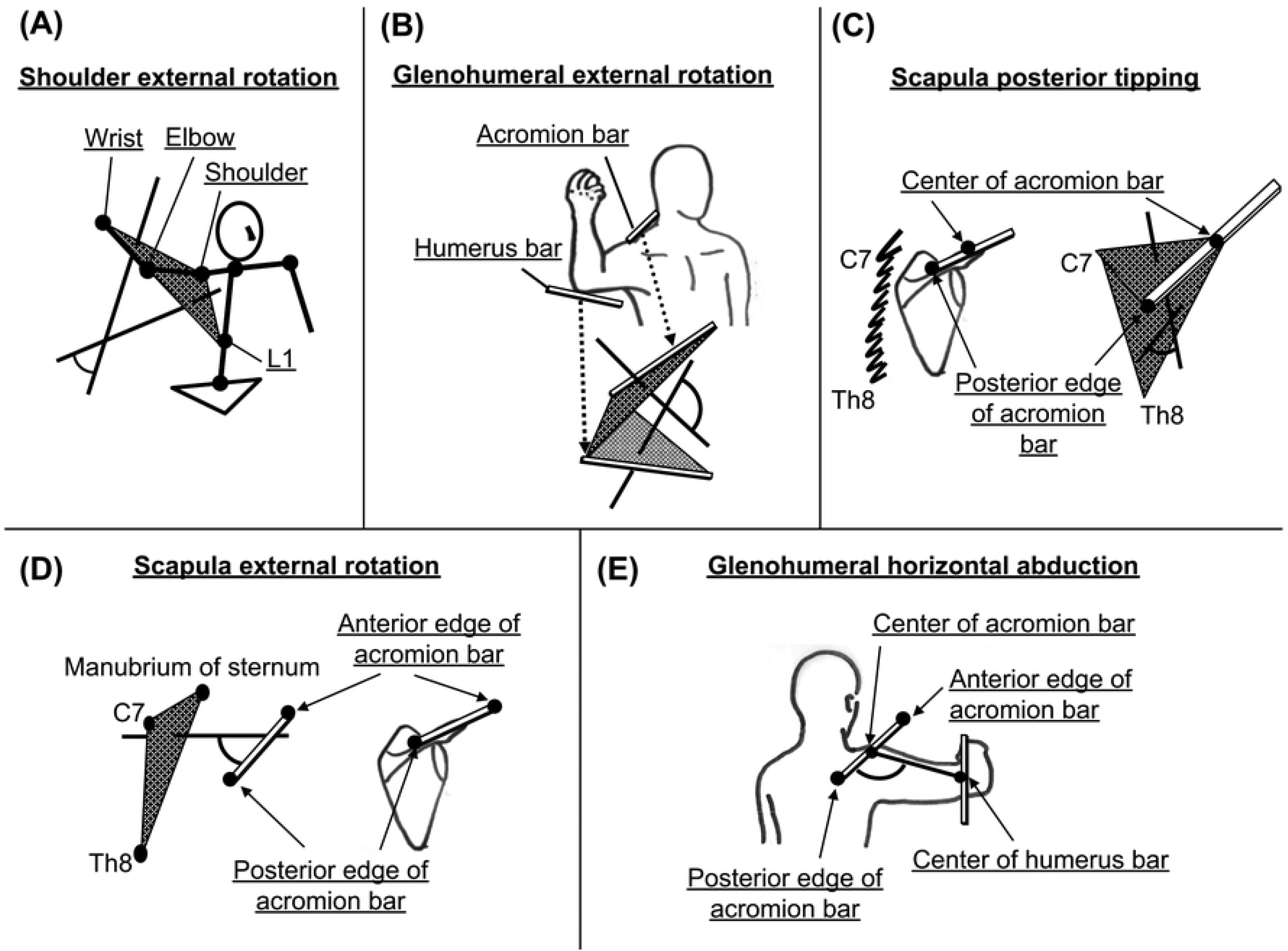
Kinematic model used for the angle calculation. (A) shoulder external rotation, (B) glenohumeral external rotation, (C) scapula posterior tipping, (D) scapula external rotation, and (E) glenohumeral horizontal abduction.

### Statistical analysis

All calculations were performed using SPSS (version 23.0, SPSS, Chicago, IL, USA). A two-way repeated measures analysis of variance (ANOVA) was used to examine the main effects of the injury and their interaction on each angle for the injury-prone group and the control group at three time points: foot contact, shoulder MER, and ball release. If the interaction was found to be significant, a post-hoc analysis was used the Bonferroni test was performed. The alpha level was considered significant when *p* < 0.05. Partial eta squared measure (η_p_^2^) was used to calculate effect sizes for ANOVA. Effect size of η_p_^2^ for the ANCOVA test was also calculated. An effect of ηp2 of ≥0.01 indicates a small effect, of ≥0.059 a medium effect, and of ≥0.138 a large effect [21].

The means and standard deviations of the amount of change in each angle during the cocking and acceleration phases were calculated. A unpaired t-test was performed to determine significant differences between groups, and values were expressed with a 95% confidence interval (CI). An alpha level was considered significant when *p* < 0.05. Cohen’s d effect sizes were used to calculate effect size for between group differences. The effect size was interpreted as small = (0.20), medium = (0.50), and large = (0.80) [21].

## Results

There was no significant difference in mean age and height between the two groups, but there was a significant difference in body mass between the two groups (Table 1).

**Table 1.** Participant Characteristics.

|  | **Injury-prone**<br>**group** | **Control group** | **t** | **p** | **d** |
| --- | --- | --- | --- | --- | --- |
|  | **(n = 15)** | **(n = 15)** |  |  |  |
| **Age, y** | 20.7 ± 1.4 | 20.9 ± 1.1 | .311 | .758 | .117 |
| **Height, cm** | 180.1 ± 6.5 | 177.1 ± 6.6 | 1.227 | .23 | .464 |
| **Mass, kg** | 78.9 ± 5.4 | 72.3 ± 6.7 | 2.968 | .006 | 1.122 |

Fig 3 shows the time curves of each joint angle. The two-way repeated measures ANOVA for scapular ER showed that the main effect of injury was statistically significant (*p* = 0.018; *F* = 6.304; *η_p_^2^* = 0.184), and the interaction effect of injury and phase was significant (*p* = 0.04; *F* = 3.485; *η_p_^2^* = 0.111) (Table 2). Post-hoc analysis of the injury × phase interaction effect showed a statistically significant differences in MER (*p* = 0.001; *F* = 13.100; *η_p_^2^* = 0.319) and ball release (*p* = 0.042; *F* = 4.560; *η_p_^2^* = 0.140) (Table 2). In addition, a significant main effect of injury was found in the assessment of the glenohumeral horizontal abduction (*p* = 0.006; *F* = 8.769; *η_p_^2^* = 0.238), with no significant interaction between injury and phase (*p* = 0.919; *F* = 0.067; *η_p_^2^* = 0.

**Figure 3.**
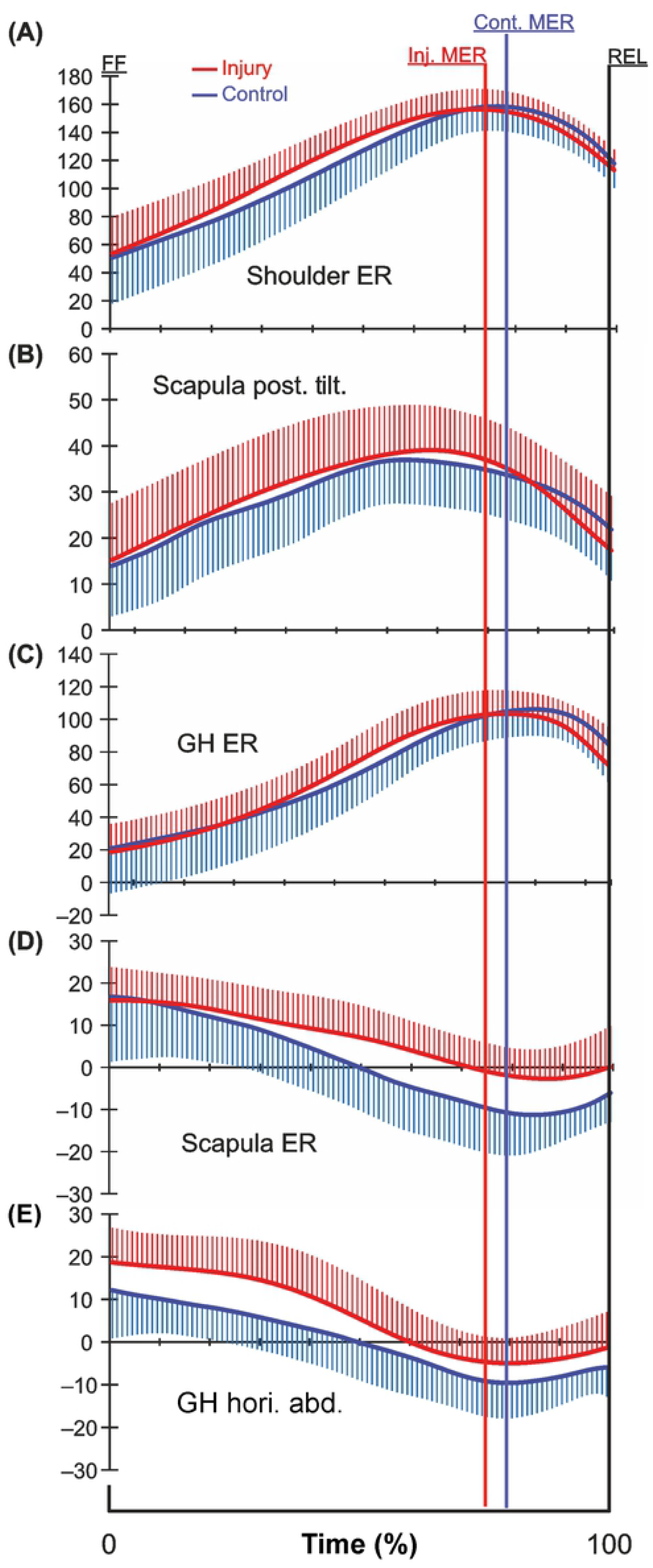
The time curve of the mean (±SD) of joint angle. (A) shoulder external rotation (ER) angle, (B) scapular posterior tilting (Post. Tilt) angle, (C) glenohumeral external rotation (GH ER) angle, (D) scapular external rotation (ER) angle, and (E) glenohumeral horizontal abduction (GH Hori. Abd.) in the throwing phase. FC, stride foot contact; MER, maximum external rotation; REL, ball release.

**Table 2.**
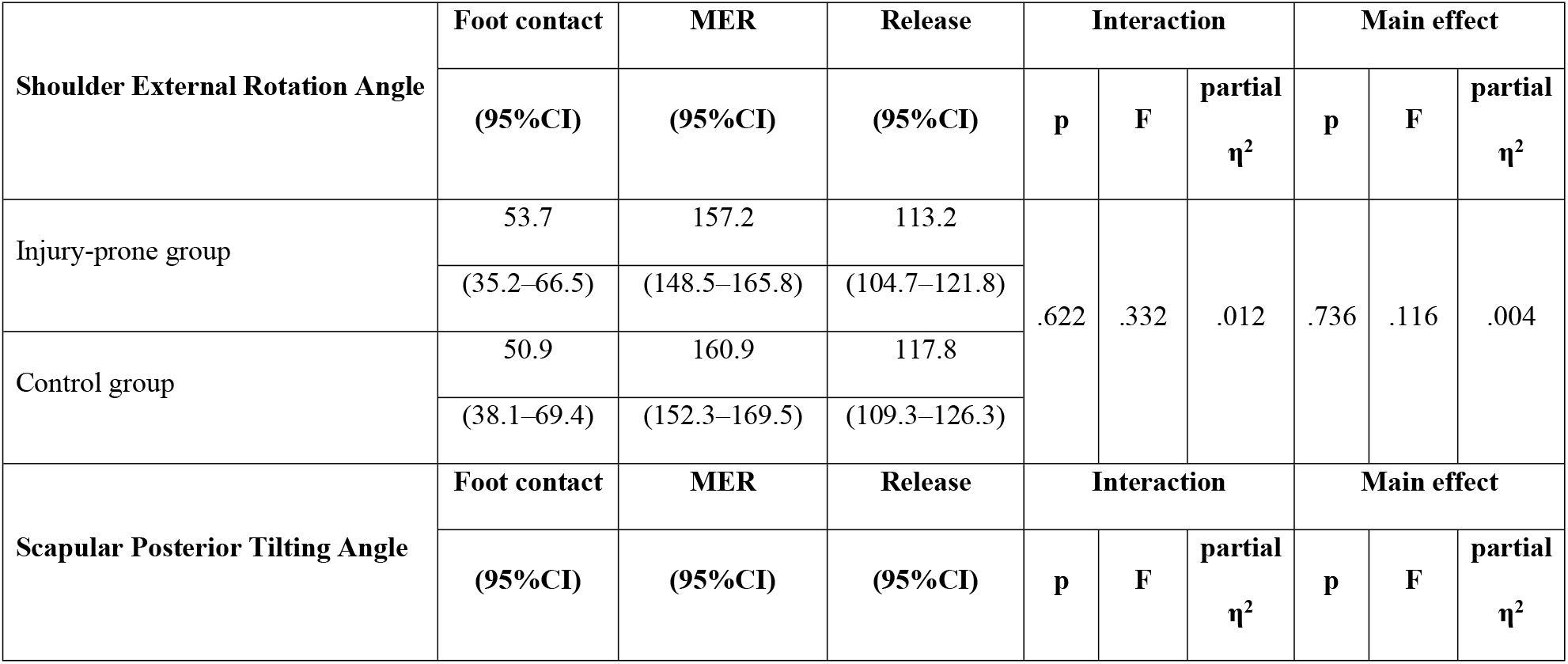
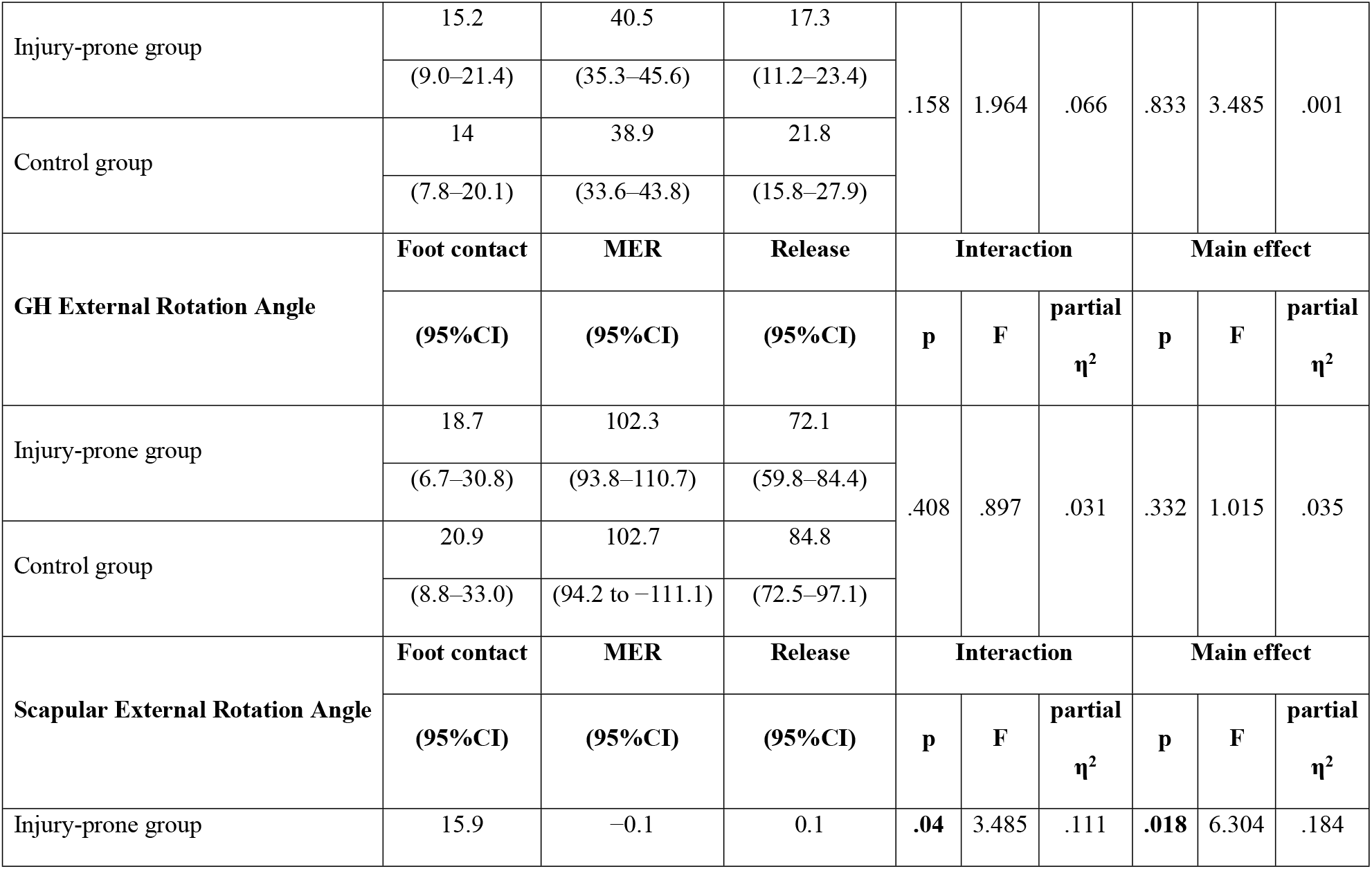
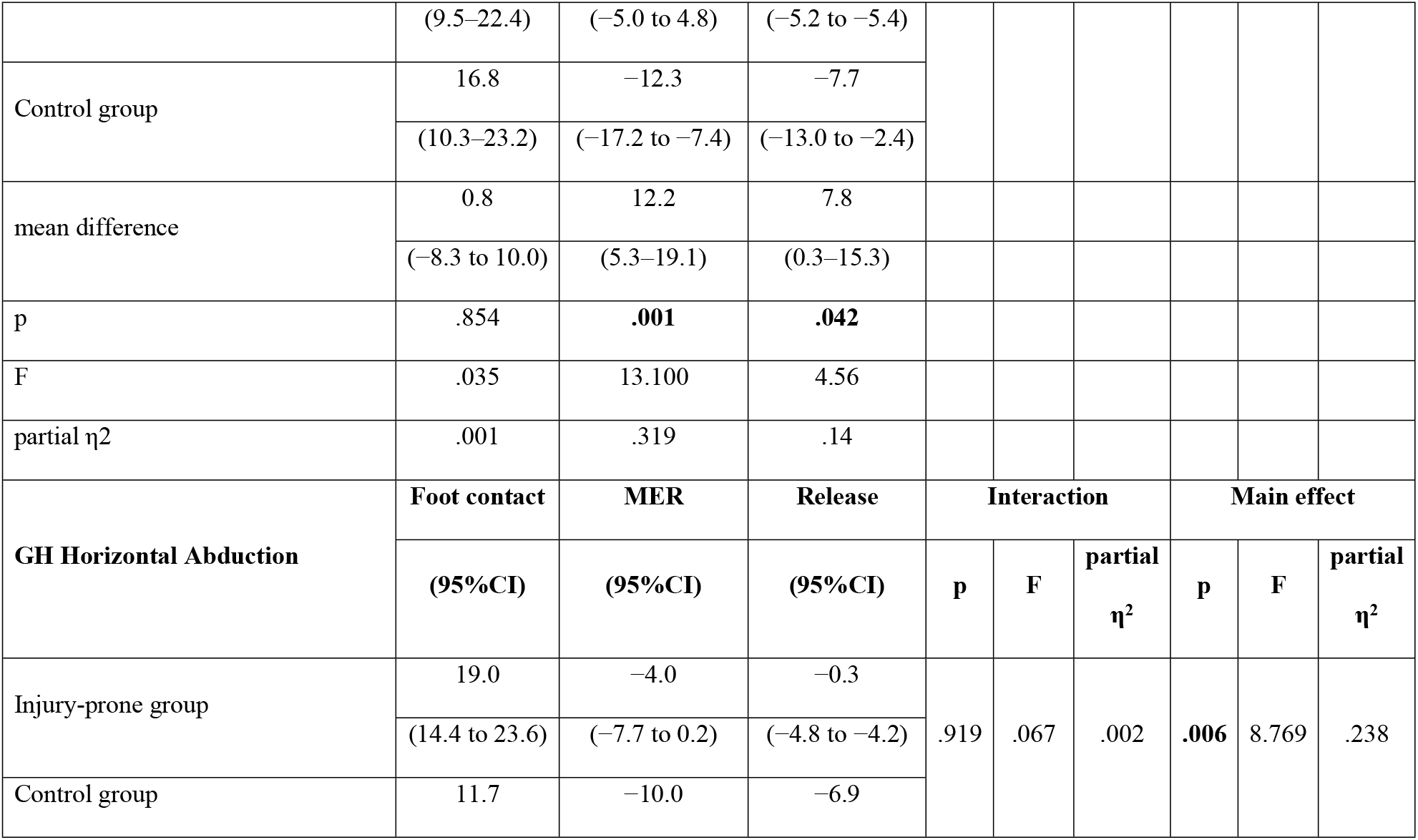
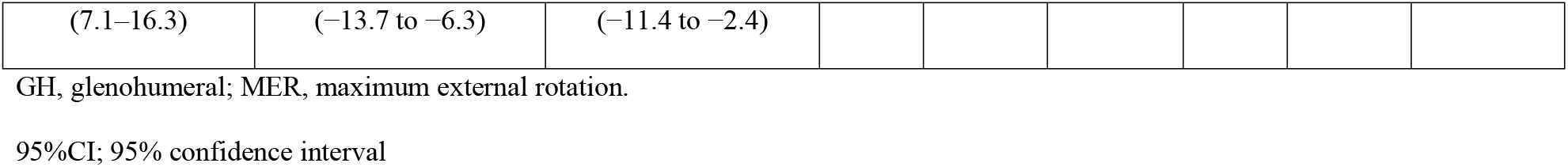
Comparison of joint angle for each phase between the injury-prone group and control group.

| Shoulder External Rotation Angle | Foot contact | MER | Release | Interaction |  |  | Main effect |  |  |
| --- | --- | --- | --- | --- | --- | --- | --- | --- | --- |
|  | (95%CI) | (95%CI) | (95%CI) | p | F | partial<br>η² | p | F | partial<br>η² |
| Injury-prone group | 53.7 | 157.2 | 113.2 | .622 | .332 | .012 | .736 | .116 | .004 |
|  | (35.2–66.5) | (148.5–165.8) | (104.7–121.8) |  |  |  |  |  |  |
| Control group | 50.9 | 160.9 | 117.8 |  |  |  |  |  |  |
|  | (38.1–69.4) | (152.3–169.5) | (109.3–126.3) |  |  |  |  |  |  |
| Scapular Posterior Tilting Angle | Foot contact | MER | Release | Interaction |  |  | Main effect |  |  |
|  | (95%CI) | (95%CI) | (95%CI) | p | F | partial<br>η² | p | F | partial<br>η² |
| Injury-prone group | 15.2 | 40.5 | 17.3 | .158 | 1.964 | .066 | .833 | 3.485 | .001 |
|  | (9.0–21.4) | (35.3–45.6) | (11.2–23.4) |  |  |  |  |  |  |
| Control group | 14 | 38.9 | 21.8 |  |  |  |  |  |  |
|  | (7.8–20.1) | (33.6–43.8) | (15.8–27.9) |  |  |  |  |  |  |
| GH External Rotation Angle | Foot contact | MER | Release | Interaction |  |  | Main effect |  |  |
|  | (95%CI) | (95%CI) | (95%CI) | p | F | partial<br>η² | p | F | partial<br>η² |
| Injury-prone group | 18.7 | 102.3 | 72.1 | .408 | .897 | .031 | .332 | 1.015 | .035 |
|  | (6.7–30.8) | (93.8–110.7) | (59.8–84.4) |  |  |  |  |  |  |
| Control group | 20.9 | 102.7 | 84.8 |  |  |  |  |  |  |
|  | (8.8–33.0) | (94.2 to –111.1) | (72.5–97.1) |  |  |  |  |  |  |
| Scapular External Rotation Angle | Foot contact | MER | Release | Interaction |  |  | Main effect |  |  |
|  | (95%CI) | (95%CI) | (95%CI) | p | F | partial<br>η² | p | F | partial<br>η² |
| Injury-prone group | 15.9 | –0.1 | 0.1 | .04 | 3.485 | .111 | .018 | 6.304 | .184 |
|  | (9.5–22.4) | (–5.0 to 4.8) | (–5.2 to –5.4) |  |  |  |  |  |  |
| Control group | 16.8 | –12.3 | –7.7 |  |  |  |  |  |  |
|  | (10.3–23.2) | (–17.2 to –7.4) | (–13.0 to –2.4) |  |  |  |  |  |  |
| mean difference | 0.8 | 12.2 | 7.8 |  |  |  |  |  |  |
|  | (–8.3 to 10.0) | (5.3–19.1) | (0.3–15.3) |  |  |  |  |  |  |
| p | .854 | **.001** | **.042** |  |  |  |  |  |  |
| F | .035 | 13.100 | 4.56 |  |  |  |  |  |  |
| partial $\eta^2$ | .001 | .319 | .14 | | | | | | |
| **GH Horizontal Abduction** | **Foot contact** | **MER** | **Release** | **Interaction** |  |  | **Main effect** |  |  |
|  | **(95%CI)** | **(95%CI)** | **(95%CI)** | **p** | **F** | **partial**<br>**$\eta^2$** | **p** | **F** | **partial**<br>**$\eta^2$** |
| Injury-prone group | 19.0 | –4.0 | –0.3 |  |  |  |  |  |  |
|  | (14.4 to 23.6) | (–7.7 to 0.2) | (–4.8 to –4.2) | .919 | .067 | .002 | **.006** | 8.769 | .238 |
| Control group | 11.7 | –10.0 | –6.9 |  |  |  |  |  |  |

|  |  |  |
| --- | --- | --- |
| (7.1–16.3) | (–13.7 to –6.3) | (–11.4 to –2.4) |

GH, glenohumeral; MER, maximum external rotation.
95%CI; 95% confidence interval

In the cocking phase, the amount of angular change in the scapular ER was significantly smaller in the injury-prone group than in the control group (mean difference, −13.0; 95% CI, −25.2 to −0.9; *p* = 0.037; *d* = 0.855) (Table 3), meaning that the injury-prone group had less internal rotation (IR) of the scapula during the cocking and acceleration phases than the control group.

**Table 3.** Comparison of the amount of joint angle change during each phase between the injury group and control group.

|  | Injury group | Control group | Mean difference (95%CI) | t | p | d |
| --- | --- | --- | --- | --- | --- | --- |
| Angle change amount during cocking phase |  |  |  |  |  |  |
| Shoulder External Rotation | 103.4 ± 27.0 | 110.0 ± 34.9 | 6.6<br>(-16.8 to 29.9) | .578 | .568 | .219 |
| Scapular Posterior Tilting | 25.2 ± 8.8 | 24.8 ± 8.5 | 0.5<br>(-6.0 to 7.0) | .164 | .871 | .062 |
| GH External Rotation | 83.5 ± 17.3 | 81.8 ± 31.2 | 1.7<br>(-17.1 to 20.6) | .188 | .852 | .071 |
| Scapular External Rotation | -16.0 ± 6.7 | -29.0 ± 21.3 | -13.0<br>(-25.2 to -0.9) | -2.261 | **.037** | .855 |
| GH Horizontal Abduction | -23.0 ± 7.5 | -21.7 ± 12.2 | 1.2<br>(-6.4 to 8.9) | .334 | .741 | .126 |
| **Angle change amount during acceleration phase** |  |  |  |  |  |  |
| Shoulder External Rotation | -44.0 ± 11.9 | -43.0 ± 15.5 | 0.9<br>(-9.4 to 11.3) | .181 | .858 | .068 |
| Scapular Posterior Tilting | -23.2 ± 7.3 | -16.8 ± 9.6 | 6.3<br>(-0.0 to 12.7) | 2.037 | .051 | .77 |
| GH External Rotation | $-30.2 \pm 26.3$ | $-17.9 \pm 24.8$ | 12.3<br>(-6.8 to 31.4) | 1.320 | .198 | .499 |
| Scapular External Rotation | $0.2 \pm 10.2$ | $4.5 \pm 14.6$ | 4.4<br>(-5.1 to 13.8) | .946 | .352 | .358 |
| GH Horizontal Abduction | $3.7 \pm 8.2$ | $3.1 \pm 6.8$ | 0.6<br>(-6.2 to 5.1) | .206 | .838 | .078 |

GH, glenohumeral; MER, maximum external rotation

## Discussion

The approximate trend of the change in the angle of horizontal shoulder abduction seen in this study was similar to the basic findings of the authors’ previous studies [22,23]. Especially, this current investigation indicates the kinematic differences of the scapular and glenohumeral movement on the horizontal plane between the injury-prone pitchers and the healthy controls.

The results of this study showed that the horizontal abduction of the glenohumeral joint during the cocking and acceleration phases of the injury-prone group was larger than that of the control group, suggesting that hyperangulation may be a risk factor for pitching-related shoulder injuries [16–18]. Furthermore, it was shown that the glenohumeral joint in the injury-prone group may not have been in a sufficient horizontal adduction position even at the MER. We speculate that in baseball pitchers with such kinematics, various factors such as pitching-related fatigue and shoulder dysfunction may further lead to increased horizontal abduction of the glenohumeral joint. Therefore, baseball pitchers who do not have proper scapular positioning during the cocking phases of pitching are considered to be at risk for injury on a daily basis.

The horizontal abduction angle of glenohumeral joint in the release motion was also significantly larger in the injury-prone baseball pitchers. Previous studies on kinematic analysis have reported that the anterior shear force at the ball release increases as the shoulder horizontal abduction angle increases [24]. Therefore, it is possible that the injury-prone group in this study also had repeated pitching-related injuries due to stress from the hyper-angulation, as has been presented in previous studies [12].

Furthermore, there were differences in the movement of the scapula in the horizontal plane between the injured group and the healthy control group. The results showed that there was no significant difference in the scapularr ER angle at the initial contact of the foot between the two groups. However, the injury-prone group had a significantly greater scapularr ER angle at MER than the control group. In addition, the amount of IR movement of the scapula during cocking was also significantly smaller in the injury-prone group than in the control group. In addition, in the injury-prone group, the IR movement of the scapula was significantly smaller during cocking phase, suggesting that inadequate IR of the scapula during the cocking phase may be associated with pitching-related shoulder injuries.

Scapula dyskinesis is considered to be a feature of pitching shoulder injuries [6,25]. It has been suggested that baseball players with pitching injuries may have the scapula in an internally rotated position and may not be able to adequately externally rotate the scapula during the cocking phase [1,6,25], leading to greater horizontal abduction of glenohumeral joint during the subsequent phase. However, the results of our study provide new insights into the relationship between the coordination of scapula and glenohumeral joint and the injury risk. In the present study, there was no significant difference in the scapular ER angle at foot contact, the starting point of the cocking phase, between the two groups. Instead, a difference was found in the subsequent scapular IR movement, suggesting that inadequate IR of the scapula during cocking and acceleration phases may increase the injury risk rather than the SICK scapular alignment.

Based on the flow of the kinetic chain, serratus anterior muscle play an important role in the IR movement of the scapular during cocking and acceleration phases, and therefore, it is important to improve the function of the serratus anterior for the prevention of pitching injuries [25]. Dysfunction of the serratus anterior muscle alters the position of the IR direction and raises the medial edge of the scapula, resulting in the so-called scapular wing [6,26]. The decreased IR movement of the scapula observed in the injury-prone group in this study may be due in part to dysfunction of the serratus anterior muscle, and maintaining and improving the function of the serratus anterior muscle may be crucial for preventing pitching-related injuries.

Scapular motion during the pitching motion also should be examined from the perspective of the kinetic chain. In the kinetic chain of pitching, the scapula is responsible for transferring rotational energy from the trunk to the upper extremities [1,25], but it can also transfer energy from distal to proximal [27]. In order for the scapula to move adequately during pitching motion and for the kinetic chain to be effective, proper positioning of the upper extremities at foot contact plays an important role. Kreighbaum et al. stated that the inertial resistance of the distal segment “stays in place” while its proximal end is pulled forward by the distal end of the proximal segment. Conversely, a force of the same magnitude due to inertial resistance is applied to the proximal segment in the opposite direction. Applying this concept to the pitching motion, if the horizontal abduction angle is already large at the time of foot contact, the rearward inertia force due to trunk rotation will be large. Simultaneously, force applied in the ER direction of the scapula will be large, resulting in the smaller IR of the scapula during cocking in the injury-prone group than in the control group. Consequently, the kinetic chain between the scapula and the glenohumeral joint, indicated by the “lagging back phenomenon,” [28] may not have worked efficiently, and the horizontal abduction angle of the glenohumeral joint may have remained larger than that of the control group in subsequent phase. From the perspective of the kinetic chain of the throwing motion, it is necessary to focus on the take-back motion during foot contact when pitching in order to prevent injuries.

In this study, there were no significant group differences in both posterior scapularr tilt and glenohumeral ER motion in all phases. It is well known that most of the pitching injuries occur in the phase close to MER. Fleisig et al. reported that an IR torque of 67 N-m and an anterior shear force of 310 N are applied to the shoulder just before MER in the pitching motion [29]. Sabick et al also reported that the peak humeral axial torque reached a mean value of 92 ± 16 N·m near the point of MER [30]. These previous findings indicate ER during the pitching motion places a great deal of stress on the glenohumeral joint and can be a major cause of pitching injury. However, in the present study, there was no difference in the magnitude of the MER angle, but there was a slight difference in the timing of the MER between the two groups. Whiteley stated that the stress on the glenohumeral joint varies greatly depending on the “timing of horizontal abduction and external rotation,” and when the glenohumeral ER is forced in the horizontal abduction position, the risk of pitching-related shoulder injuries may be increased [31]. The relative positions of horizontal abduction and external rotation of the GH during pitching need to be further investigated to determine how they affect stress on the shoulder.

The limitation of this study is that it did not analyze joint function. Therefore, it is unclear whether the differences in scapular motion are due to joint dysfunction or kinetic chain problems. Also, the validity of this analysis method needs to be further investigated. Due to ethical issues and technical limitations, we cannot compare our method with other methods such as radiography to verify its accuracy. Instead, our preliminary study [32] validated the current method by comparing it to methods that uses electromagnetic sensors. In addition, the shoulder joint ER and the horizontal abduction angle of the shoulder during pitching obtained by the current method were quite consistent with those obtained by a three-dimensional motion analysis system equipped with 10 cameras [24] and an electromagnetic tracking device [33]. In addition, our method has the technical advantage that the participants can throw with maximum effort since there are no restrictions on the throwing motion due to the wearing of the sensor. Another advantage is that the movement of the scapula and humerus can be visualized indirectly through a bar attached to the body, as shown in video 1.

This study demonstrated that there were differences between injury-prone and healthy counterparts in the movement of the scapula and glenohumeral joint during the pitching motion. Three-dimensional analysis revealed that the injury-prone group had a significantly greater horizontal abduction angle than the control group. Furthermore, there was a difference in the movement of the scapula on the horizontal plane between the groups; the injury-prone group had a greater ER angle of the scapula than the control group in MER and ball release, but there was no difference in foot contact between the groups. Also, the amount of decrease in scapular ER angle during the cocking phase was significantly smaller in the injury-prone group than in the control group, suggesting that injured baseball pitchers do not have sufficient control over scapular IR during the cocking and acceleration phases. In order to establish effective preventive measures against pitching-related shoulder injuries, further research is needed to determine whether the biomechanical characteristics of the scapula and glenohumeral joint in injury-prone pitchers are the result of dysfunction of the scapula or a problem in the kinetic chain during pitching.

## Acknowledgments

The authors would like to thank Enago (www.enago.jp) for the English language review.

## Author Contributions

- Conceived and designed the analysis: Koji Miyashita, Sentaro Koshida, Taro Koyama, Kenicihro Ota, Yusuke Tani, Ryoji Okamune
- Data collection: Koji Miyashita, Sentaro Koshida, Taro Koyama, Kenicihro Ota, Yusuke Tani, Ryoji Okamune
- Contributed data/analysis tools: Koji Miyashita, Sentaro Koshida, Taro Koyama, Kenicihro Ota, Yusuke Tani, Ryoji Okamune
- Performed the analysis: Koji Miyashita, Sentaro Koshida, Taro Koyama, Kenicihro Ota, Yusuke Tani, Ryoji Okamune
- Wrote the paper: Koji Miyashita, Sentaro Koshida, Taro Koyama, Kenicihro Ota, Yusuke Tani, Ryoji Okamune

